# Laboratory strains of *Aedes aegypti* are Competent to Brazilian Zika virus

**DOI:** 10.1101/071654

**Authors:** André Luis Costa-da-Silva, Rafaella Sayuri Ioshino, Helena Rocha Corrêa Araújo, Bianca Burini Kojin, Paolo Marinho de Andrade Zanotto, Danielle Bruna Leal Oliveira, Stella Rezende Melo, Edison Luiz Durigon, Margareth Lara Capurro

## Abstract

Since the Zika outbreaks are unprecedented human threat in relation to congenital malformations and neurological/autoimmune complications as well as its high potential to spread in regions presenting the vectors, improvements in mosquito control is a top priority. Thus, *Aedes aegypti* laboratory strains will be fundamental to support studies in different research fields implicated on Zika-mosquito interactions which are the basis for the development of innovative control methods. In this sense, we determined the main infection aspects of the Brazilian Zika strain in reference *Aedes aegypti* laboratory mosquitoes.

We orally exposed Rockefeller, Higgs and Rexville mosquitoes to a Brazilian ZIKV (ZIKV^BR^) and qRT-PCR was applied to determine the infection and dissemination rates, and viral levels in mosquito tissues as well as in the saliva. The ZIKV^BR^ kinetics was monitored during the infection in Rockefeller mosquitoes. Rockefeller strain was the most susceptible at 7 days post-infection but all strains presented similar infection levels at 14 days post-infection. Although variations in the saliva detection rates were observed, we confirmed that ZIKV^BR^ was present in saliva from Rockefeller, Higgs and Rexville females at detectable levels at 14 days post-infection. The ZIKV^BR^ kinetics in Rockefeller mosquitoes showed that the virus could be detected in the heads at 4 days post-infection but was more consistently detected late in infection. The viral levels peaked at 11 days post-infection in the mosquito bodies, remaining stable until 14 days post-infection, in contrast to the heads, where the mean viral levels only peaked at 14 days post-infection.

Our study presents the first evaluation on how Brazilian Zika virus behaves in reference *Aedes aegypti* strains and shed light on how the infection evolves over time. Vector competence and basic hallmarks of the ZIKV^BR^ development were revealed in laboratory mosquitoes. This study provides additional information to accelerate studies focusing on ZIKV-mosquito interactions.

## Introduction

Currently, the world is facing a new outbreak of the emerging Zika virus (ZIKV) [1]. Its association with neurological and autoimmune complications as well as infants born with microcephaly [2, 3] has caused a global healthcare crisis. Due to the severe situation, it was launched a document containing an operation plan to help affected countries to establish a strategy to control the disease and improvements in vector control were highlighted as priorities [4].

The outcome of vector infection will rely on the specific interactions between the mosquito and virus genotypes. Therefore, better understanding of the mosquito vectors-ZIKV interactions is the basis to generate the development of innovative strategies that can be added to the arsenal in the combat of ZIKV. Recently studies have reported significant differences in susceptibility for ZIKV infection between wild mosquito populations of *Aedes aegypti*, *Aedes albopictus* and other *Aedes* species [5–7]. These vector competence studies focusing main vector species as well as variations in susceptibility of different populations from the same species are primordial to delineate improved control programs, prioritizing competent populations.

Although wild populations of the main vector, *Ae. aegypti*, represent the natural dynamics of the ZIKV infection process in the invertebrate host, the determination of the vector competence of different laboratory strains, which are well-adapted in captivity and readily available for experiments, is also essential to support basic and applied studies in different research fields related to vector-virus interactions. Moreover, some strains are also known to be standard in many laboratories in the world and because the experimental reproducibility is more robust than with field populations, they are used as reference mosquito strains [8]. Important advances on mosquito immune responses to dengue virus (DENV) and other human pathogens have been performed on *Ae. aegypti* laboratory strains and natural mosquito populations for the purposes of comparison [8–10] as well as for the evaluation of insecticide resistance [11, 12]. Furthermore, the characterization of the vector competence is relevant for more applied purposes. The development of transgenic mosquitoes mainly uses laboratory reference strains for transformation and the genetic background related to pathogen susceptibility is incorporated to the established lines. Thus, the vector competence of laboratory strains to ZIKV must be also considered in the context of the production and release of transgenic mosquitoes [13].

To generate basic information about the interaction between ZIKV^BR^and three important laboratory-maintained *Ae. aegypti* strains, Rockefeller, Higgs white eyes and Rexville mosquitoes were analyzed. The ROCK strain, an insecticide-susceptible strain of Caribbean origin that was established in 1930 [14], is commonly used as a reference for insecticides resistance trials and in DENV infection experiments [15, 16]. The HWE strain is an eye-pigment-deficient *Ae. aegypti*, a variant of the Rex-D strain, and it is used as the recipient for mosquitoes germ-line transformations, because the lack of pigment is a desirable phenotype for visual screening of transgenic individuals, which are often marked with a fluorescent protein expressed in the eyes [17]. The RED strain of *Ae. aegypti*, also a variant from Rex-D strain, is widely used to investigate pathogen-host susceptibility [18].

This report details the ZIKV^BR^infection, dissemination and saliva detection rates in these three mosquito strains and reveals the viral kinetics in the ROCK reference strain.

## Material and methods

### Ethics Statement

The use of human blood or its derivatives products were conducted according to the principles expressed in the Declaration of Helsinki for mosquito feeding experiments. The approval of the protocol was provided by the Institutional Review Board in Human Research (IRB) (Comissão de Ética em Pesquisa com Seres Humanos do Instituto de Ciências Biomédicas/USP-CEPSH) and National Committee for Ethics in Research (Comissão Nacional de Ética em Pesquisa – CONEP), protocol #503. Number of approval: 914.876 (CAAE 38518114.0.0000.5467).

The Brazilian Zika virus strain, named as ZIKV^BR^, was previously isolated from a Brazilian clinical case [2] and a lyophilized virus sample was gently provided by the Evandro Chagas Institute in Belém, Pará. The use of the anonymized viral samples were approved by our Institutional Review Board (IRB). The research was approved by the Ethics Committee on the Research with Humans (CEPSH-Off.011616) of the Institute of Biosciences of the University of São Paulo.

### Mosquito rearing

The experiments were performed using three *Ae. aegypti* laboratory strains: HIGGS white-eye (HWE), Rexville D (RED) and Rockefeller (ROCK). Those mosquitoes were maintained in a BSL-2 insectary facility in Institute of Biomedical Sciences from University of São Paulo. The rearing conditions were 27 ± 1°C, 75-80% relative humidity with 12/12 hour (light/dark) photoperiod. Adult mosquitoes were maintained *ad libitum* on 10% sucrose solution (w/v).

### Viral amplification and titration

The Brazilian Zika virus strain, named previously as ZIKV^BR^, was isolated from a Brazilian clinical case [2] and a lyophilized virus sample was gently provided by the Evandro Chagas Institute in Belém, Pará. ZIKV^BR^ was amplified and titrated as recently described [2]. Viral titrated aliquots (5.0 x 10^6^ plaque forming unit [pfu]/mL) of fourth subculture (T4) was provided by the ZIKV São Paulo task force.

### Mosquito Infection

Pre-mated 7-9 day old *Ae. aegypti* females were sucrose 10%-deprived for 24 hours prior blood meal. Starved females received ZIKV^BR^ infectious blood meal by using the Glytube artificial feeder [19]. Human concentrated erythrocyte was mixed with ZIKV^BR^ supernatant and inactivated human serum in a 10:10:1 proportion, respectively and the ZIKV^BR^ final titer in this mixture was 2.2 x 10^6^ pfu/mL. The infected-blood was offered to female mosquitoes for 45 minutes. Non-engorged females were removed and engorged ones were kept in plastic cups maintained with 10% sucrose solution until the collection times.

### Mosquito saliva and tissue samples

Saliva, heads and bodies from individual mosquitoes (10 each), were collected at 7 and 14 dpi for virus detection. For saliva, the forced salivation technique as used as previously described [20, 21], with modifications. Mosquitoes were CO_2_ anaesthetized, transferred to a Petri dish on ice and legs and wings were removed with forceps. A glass slide with a strip of modeling clay was used to support micropipette tips filled with 10 µL of Leibovitz’s L-15 medium (Gibco^tm^) and the proboscis of each live mosquito were inserted into the tip. Insects were allowed to salivate for 45 minutes and total volume was ejected into the 1.5 mL microtube. After salivation, heads were separated from the rest of the bodies using a McPherson-Vannas Scissors #501234 (World Precision Instruments, Sarasota, FL) and each tissue was transferred to a 1.5 mL microtube. Bodies, heads and saliva samples collected at 7 and 14 dpi were immediately frozen in dry ice and stored at -80^0^C.

### RNA extraction of mosquito samples

Total RNA from individual heads, bodies and saliva were extracted using QIAmp Viral RNA Mini Kit (Qiagen, Valencia, CA, USA) following manufacturer’s recommendations. Total RNA was eluted in 60 µL of elution buffer and kept in -80°C until qRT-PCR analyses.

### One-step qRT-PCR analysis

To detect ZIKV and to quantify viral copy numbers, the Power SYBR^®^ Green RNA-to-C_T_^TM^ *1-Step* kit (Applied Biosystems, Foster City, CA, USA) was used in qRT-PCR reactions as described by the manufacturer. Each sample was analyzed in technical duplicates in a 20 µL final volume reactions containing 5 µL of total RNA and 0.5 µM for each ZIKV 835 and ZIKV 911c primers [22]. Negative (RNAse free water) and positive controls (500 ηg of total RNA extracted from ZIKV cell culture supernatant aliquot) were used in each assay. Samples were considered positive for ZIKV only when detected in both analyses. The assays were performed in Mastercycler Realplex 2 thermocycler (Eppendorf) with the following conditions: 48°C for 30 min and 95°C for 10 min followed by 40 cycles of 95°C for 30 sec, 60°C for 30 sec and a melting curve step of 95°C for 1 min, 60°C for 30 sec and 95°C for 1 min, with a temperature ramping from 60°C to 95°C at 0.02°C/sec. Amplicon specificity was evaluated by the peak of the melting curve (79 ± 1°C) and primer pair efficiency ranged from 1.01 to 1.02.

For ZIKV copy number estimation, a standard curve was generated as described [23], with some modification. Briefly, a target plasmid containing a 76 bp ZIKV fragment amplified with ZIKV 835 and ZIKV 911c primer pairs was linearized and nine serial dilutions ranging from 10^−9^ to 10^−17^ g were used to produce a standard curve. The limit of detection was experimentally established in 23 copies (10^−16^ g dilution). The ZIKV absolute copy numbers were extrapolated from the standard curves (R^2^ ranged from 0.990 to 0.996), adjusted by back-calculation to the total RNA volume (60 µL) and were expressed as copies per tissue.

### Infection, dissemination and saliva detection rate analysis

Following the definitions proposed in [5]. The adopted infection rate (or prevalence) is determined as the proportion of mosquitoes with virus detected in body (abdomen and thorax) among tested ones. Dissemination rate corresponds to the number of mosquitoes with infected heads among infected bodies (abdomen and thorax positive). Since a method to detect viral RNA in saliva was used, the term saliva detection rate was applied in place of transmission rate (which refers to infectious particles present in saliva), but with equivalent meaning. The saliva detection rate was estimated as the proportion of mosquitoes with positive saliva among mosquitoes with disseminated infection (positive heads) and was expressed as percentages.

### Data analysis

Statistical analyses were performed in GraphPad Prism5 software package (Version 5.00) for Windows (San Diego, California, USA) and confidence intervals of 95% were defined for all analyses. Fisher’s exact test were applied in conformity with [24] to detected significant differences in ZIKV positive proportions of the same tissues (bodies or heads or saliva) at 7 or 14 dpi from different mosquito strains and to compare infection proportions during ZIKV kinetics in ROCK strain. Viral levels between different *Ae. aegypti* lines were compared by using Two-way ANOVA and Bonferroni posttests.

## Results

### ZIKV^BR^ infection prevalence and dissemination rates in orally challenged mosquito strains

To analyze and compare the infection susceptibility of *Ae. aegypti* laboratory strains to a Brazilian ZIKV, we orally exposed ROCK, HWE and RED mosquitoes with a low-passage ZIKV^BR^ strain. Viral RNA levels were quantified by qRT-PCR assay in individual mosquito bodies and heads at 7 and 14 days post-infection (dpi). These intervals are well-characterized indicators for infection establishment in the abdomen and viral dissemination to the head, respectively, during flavivirus replication progression in *Ae. aegypti* mosquitoes [25].

The three strains exhibited high viral levels in the bodies at 7 and 14 dpi, with the mean viral copy numbers fluctuating from 10^7^ at 7 dpi to 10^8^ at 14 dpi (Fig 1A). At 7 dpi, the ROCK and RED strains showed the highest body infection rates (number of positive bodies/total mosquitoes tested) (90%) compared to HWE females (70%). At 14 dpi, the body infection proportion increased in HWE samples and all the strains showed the same infection rate (90%) at this time. However, the infection rate of the heads exhibited variation in the number of positive samples among the strains at 7 dpi, with ROCK at 90%, HWE at 50% and RED at 60%. At 14 dpi, the ROCK strain remained with 90% of the tested heads infected and the HWE and RED strains had increases in the percentage of infected heads to 80% and 90%, respectively. With regard to the mean viral levels observed in the head, we observed that the RED strain had 10^5^ ZIKV copies at 7 dpi and this number increased to 10^7^ at 14 dpi, indicating an increase of 2 logs after 7 days, while the ROCK and HWE strains had infection level increases only 1 log (Fig 1B). When analyzing the dissemination rate (the number of infected heads/number of infected bodies), the ROCK strain had the highest index (100%) analyzed at any time (Fig 2).

**Fig 1.**
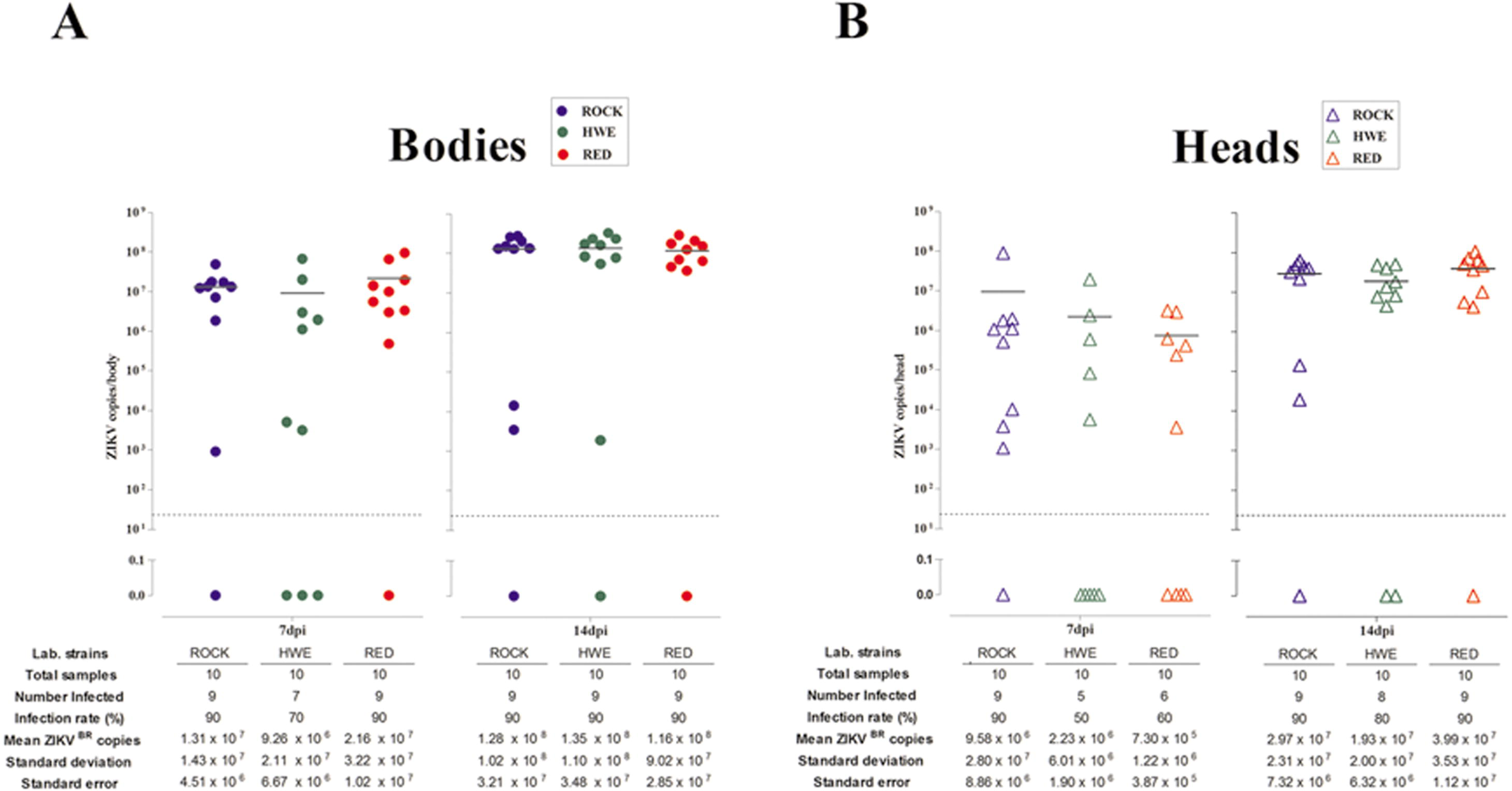
ZIKV^BR^ infection rates and viral levels in bodies and heads from *Ae. aegypti* laboratory strains. The infection rate and viral levels per tissue were individually recorded in ROCK, HWE and RED females at 7 and 14 days following a ZIKV-infected blood meal (dpi). (**A**) Viral prevalence and infection levels in the bodies. Each body sample is represented by a solid circle. (**B**) Viral prevalence and infection levels in the heads. Individual heads are indicated by open triangles. Black bars indicate the mean viral copy numbers and the dashed grey line demonstrates the detection limit. Significant differences were not observed between the bodies or heads from the three strains at 7 and 14 dpi in the infection prevalences by Fisher’s exact test (p>0.05), or in the viral infection levels by two-way ANOVA with Bonferroni post-tests (p>0.05)

**Fig 2.**
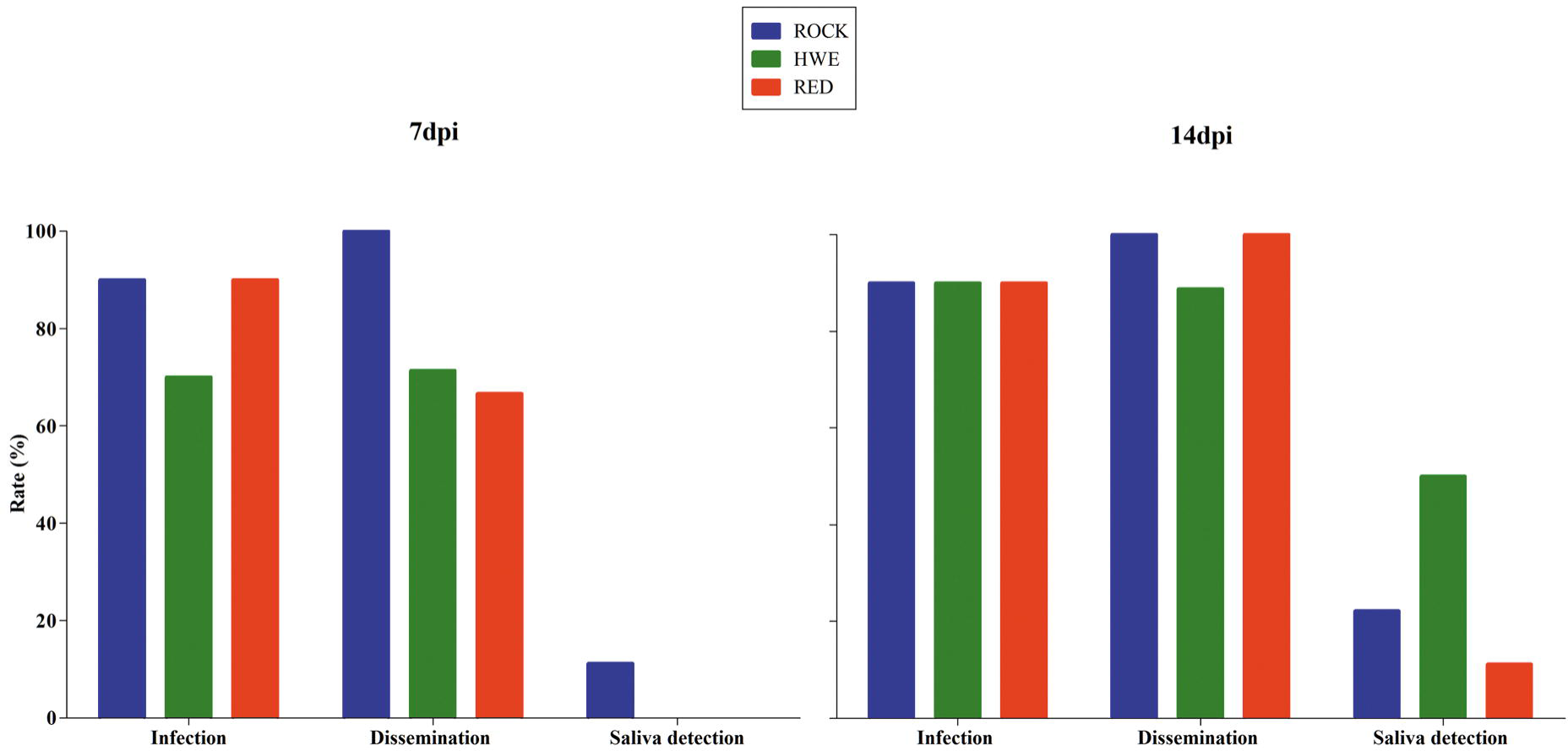
ZIKV^BR^ infection, dissemination and saliva detection rates in *Ae. aegypti* laboratory strains. The infection rate (number of infected bodies/total bodies analyzed), dissemination rate (number of infected heads/number of infected bodies) and saliva detection rate (number of infected saliva sample/number of infected heads) were estimated in females from ROCK, HWE and RED strains at 7 and 14 days following a ZIKV-infected blood meal (dpi). The results are represented by percentages.

Although strain variations in the infection rates and mean levels of ZIKV^BR^ were characterized in the bodies and heads at 7 and 14 dpi, no significant differences were observed in tissue infection prevalence (Fisher’s exact test, p>0.05) and viral intensity between the ROCK, HWE and RED samples (two-way ANOVA followed by Bonferroni post-tests, p>0.05).

### Prevalence and detection rate of ZIKV^BR^ in mosquito saliva

The presence of ZIKV^BR^ and viral level quantitation were assessed directly for each collected saliva sample using the same qRT-PCR assay. ROCK strain females showed one positive saliva sample (10%) at 7 dpi. However, no ZIKV^BR^ was detected in saliva from HWE and RED mosquitoes at this time point. In contrast, saliva samples from the three strains presented detectable viral levels at 14 dpi. HWE samples were 40% positive for ZIKV^BR^, while ROCK and RED samples were 20% and 10%, respectively (Fig 3). Accordingly, the highest ZIKV^BR^ saliva detection rate (number of positive saliva samples/total number of infected heads) was observed in the HWE strain (50%), followed by ROCK (22.22%) and RED (11.11%).

**Fig 3.**
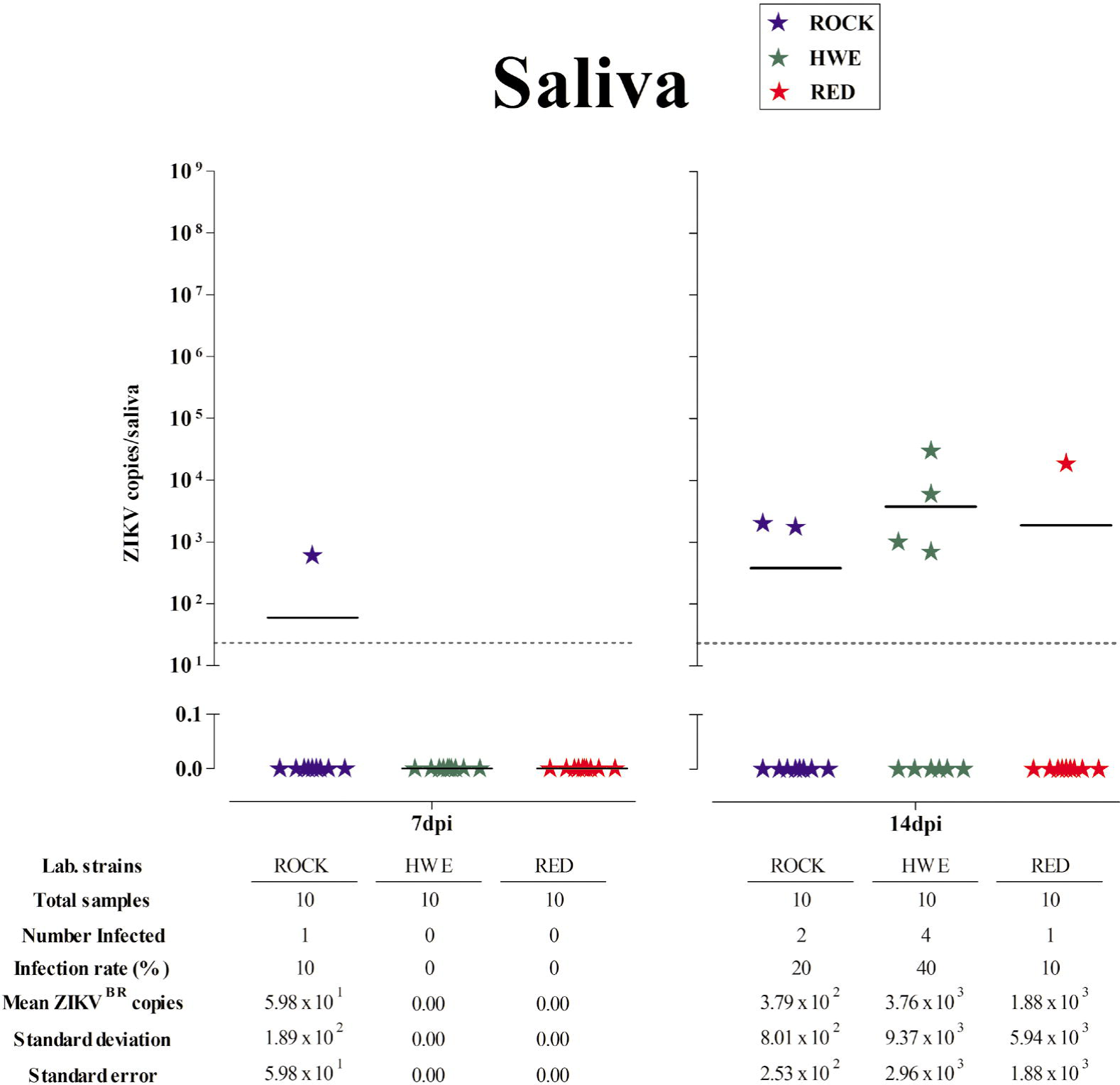
ZIKV^BR^ prevalence and viral levels in collected saliva from *Ae. aegypti* laboratory strains. The prevalence and viral levels per saliva sample were individually recorded in ROCK, HWE and RED strains at 7 and 14 days following a ZIKV-infected blood meal (dpi). Each saliva sample is represented by a solid star. Black bars indicate the mean viral copy numbers and the dashed grey line demonstrates the detection limit. Significant differences were not observed between samples from the three strains at 7 and 14 dpi in saliva infection prevalences by Fisher’s exact test (p>0.05), or in viral detection levels by two-way ANOVA and Bonferroni post-tests (p>0.05).

The ZIKV^BR^prevalence was low in the ROCK and RED saliva samples. The mean viral levels in ROCK saliva increased 1 log (10^1^ to 10^2^) between 7 and 14 dpi. In comparison with ROCK, It was observed higher mean ZIKV levels in the RED and HWE samples (10^4^) (Fig 3) at 14 dpi.

As observed in the bodies and heads, saliva samples showed strain variations related to the detection rate and the mean ZIKV^BR^ levels (Figs 2 and 3, respectively). However, no statistically significant differences were detected in virus prevalence (Fisher’s exact test, p>0.05) or levels in the saliva (two-way ANOVA followed by Bonferroni post-tests, p>0.05) between the ROCK, HWE and RED strains at 7 and 14 dpi.

### ZIKV^BR^ invasion and establishment kinetics in the ROCK strain

ZIKV^BR^infection, dissemination and saliva detection rates were higher in the ROCK strain at 7 dpi (Fig 2). This result led us to perform an independent experiment in order to determine the kinetics of ZIKV^BR^ during the establishment and dissemination of infection in females from this strain. Five time-points post-infected blood meal (1, 4, 7, 11 and 14 dpi) were used to perform a quantitative analysis of the viral levels in the individual bodies and heads (Fig 4).

**Fig 4.**
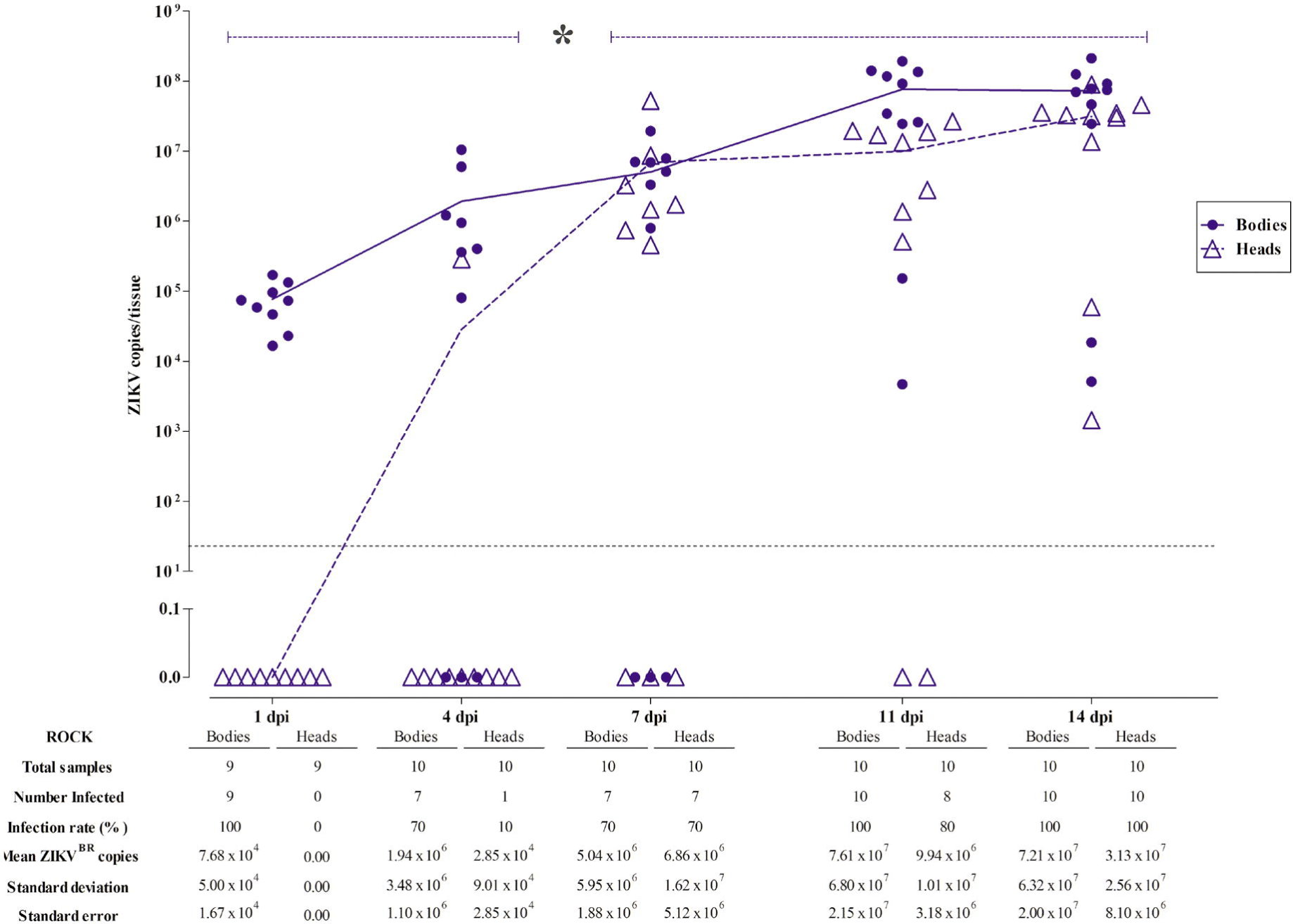
ZIKV^BR^ kinetics in bodies and heads of *Ae. aegypti* from the ROCK strain. The infection rate and viral levels per tissue were individually recorded at 1, 4, 7, 11 and 14 days following a ZIKV-infected blood meal (dpi). Solid circles and open triangles represent each body and head sample, respectively, from the ROCK females. Solid (body) and dashed (head) blue lines indicates the viral infection progression (mean levels) during the time-course experiment. Dashed blue bars indicate early (1 to 4 dpi) and late (7 to 14 dpi) infection stages in heads in which a significant difference in infection prevalence was observed between the two stages using Fisher’s exact test (*, 1 dpi x 7 dpi - p=0.0031; 1 dpi x 11 dpi - p=0.0007; 1dpi x 14 dpi - p<0.0001; 4 dpi x 7 dpi - p=0.0198; 4 dpi x 11 dpi - p=0.0055; 4 dpi x 14 dpi - p=0.0001). The dashed grey line demonstrates the detection limit.

As expected, all the body samples analyzed were positive for ZIKV^BR^ at 1 dpi and the mean viral levels was 7.68 x 10^4^ copies. These results probably reflect the viral load ingested during the oral infection of the ROCK females and ZIKV^BR^ particles still present in the infectious blood remaining in the blood bolus inside the midgut. Not surprisingly, none of the heads showed detectable levels of ZIKV in this period.

The detected viral copies increased gradually in the bodies and heads over the experimental time course. At 4 dpi, when the blood had been completely digested, the prevalence was 70% in the bodies and 10% in the heads, indicating that in some ROCK females, ZIKV^BR^ infection can evolve more rapidly, reaching the head and producing high infection intensity in the nervous tissues (10^5^) over a short period after infection (Fig 4).

The amount of infected bodies remained at the same level (70%) at 7 dpi, however, the number of virus-infected heads increased to 70% and the mean viral levels were enhanced by approximately 2 logs. ZIKV^BR^ was detected in the bodies of all analyzed mosquito samples at 11 and 14 dpi. Furthermore, a viral level peak was reached at 11 dpi, and an infection plateau was maintained in the body tissues between the last two sampling times (Fig 4).

The ZIKV^BR^ prevalence was 80% in the heads at 11 dpi and 100% at 14 dpi. The mean viral levels increased during the 14 days of infection from 0.0 to 3.13 X 10^7^ copies in the heads of mosquitoes fed with a viral titer of 2.27 x 10^6^ pfu/mL (final concentration). The infection rates of the ROCK head samples varied significantly between the early (1 dpi and 4 dpi) and late time points of infection (7, 11 and 14 dpi) (Fisher’s exact test, p<0.05).

## Discussion

Basic knowledge on the interactions ZIKV with its vectors is one of the priorities in order to create a solid scientific foundation supporting traditional and innovative methods to face the Zika challenge [26]. The literature in this field is expanding with recent studies uncovering the main species naturally infected with ZIKV [27–29] and characterizing the viral susceptibility of the natural populations in regions with the potential for urban transmission [5, 30]. As important as these studies including wild or recently colonized mosquitoes are, well-adapted laboratory vector strains will provide a consistent basis for reliable cellular and molecular studies of the virus-mosquito interaction, in which execution feasibility and reproducibility are essential.

Recently, the laboratory-adapted mosquito strains, HWE and Orlando (ORL), were used to describe the infection pattern of chikungunya virus (CHIKV). Both mosquito strains were susceptible to the CHIKV, and viral particles were detected in the saliva only two days after an infectious blood meal. The CHIKV infection pattern in midguts and dissemination rate were significantly lower for the ORL in comparison to the HWE strain until 3 dpi, although the HWE and ORL mosquitoes showed similar rates of virus in the saliva (60 and 65%, respectively) at 7 dpi [31].

Our study also found variations between laboratory-adapted strains during the ZIKV infection. Although *Ae. aegypti* infection dynamics is more rapid for the CHIKV (an alphavirus member from the Togaviridae family) than the ZIKV (a flavivirus from the Flaviviridae family), the HWE strain demonstrates lower saliva prevalence in comparison to the ROCK strain in early sampling time during ZIKV^BR^ infection. The same pattern was observed in the HWE mosquitoes in relation to the ORL strain when exposed to the CHIKV [31].

CHIKV prevalence into saliva of the HWE increases from 20% at 2 dpi to 60% at 7 dpi, differing from the ORL strain (55% at 2 dpi to 65% at 7 dpi)[31]. Our study also demonstrates that the increase of the ZIKV^BR^ saliva detection rate was more pronounced in the HWE infected mosquitoes (0% at 7 dpi to 50% at 14 dpi) in relation to the RED (0% at 7 dpi to 11.1% at 14 dpi) and ROCK (11.1% at 7 dpi to 22.2% at 14 dpi). This result is surprising since the HWE has the lowest infection rate among the strains at 7 dpi while the ROCK mosquitoes showed ZIKV^BR^ susceptibility that results in faster infection establishment and dissemination. More interestingly, the HWE strain showed the highest ZIKV^BR^ load in the saliva at late infection stage and a similar result was demonstrated for the HWE mosquitoes infected with CHIKV [31].

American populations of *Ae. aegypti* were orally exposed to an Asian genotype of ZIKV and viral infection and dissemination were observed in the early days post-infection. Although the infection rates were high, dissemination and transmission rates were comparatively low [5]. The infection and saliva detection rates were similar to our results, but we found high dissemination rates in all *Ae. aegypti* strains tested.

Consistent with our findings with ZIKV^BR^ infection in *Ae. aegypti* laboratory strains, other studies found the same high susceptibility in wild mosquito populations infected with different ZIKV strains, highlighting that the reference strains can mimic the infection pattern of wild population [5, 32].

The present work adopted qRT-PCR as a rapid and efficient method to characterize vector competence [7] and to precisely measure the viral levels during the infection process [24, 33]. Studies have shown a consistent correlation between viral RNA levels and infectious viral particles of different flaviviridae [34, 35]. Based on ZIKV^BR^ genome amplification, we measured the saliva detection rates to verify the ZIKV^BR^ competence of three *Ae. aegypti* laboratory strains. The detection of virus RNA in the mosquito saliva indicates that salivary gland infection and escape barriers were overcome and implies that *Ae. aegypti* mosquitoes from the ROCK, HWE and RED strains are competent to ZIKV^BR^.

## Conclusions

The results from our study confirm that ROCK, HWE and RED laboratory strains not only sustain the development of the Brazilian Zika virus but are also competent for virus transmission. These findings provide useful comparisons for future researches and will dictate the strains that suits best for desired experiments. In this sense, this knowledge is fundamental for Zika-invertebrate host studies, especially because we determined the main infection aspects of the ZIKV^BR^ strain in reference *Aedes aegypti* laboratory mosquitoes. This knowledge is the first step to support the researches aiming to understand ZIKV-vector biology focusing innovative solutions on vector control.

## Acknowledgements

We thanks Dr. Pedro Vasconcelos from Evandro Chagas Institute IEC in Belém for providing the lyophilized ZIKV isolate, Carla Torres Braconi for technical advices and Isabel Cristina dos Santos Marques and Ediane Saraiva Fernandes for technical assistance.

